# A transformer model for predicting cognitive impairment from sleep

**DOI:** 10.1101/2022.07.17.500351

**Authors:** Tzu-An Song, Masoud Malekzadeh, Richa Saxena, Shaun M. Purcell, Joyita Dutta

**Affiliations:** University of Massachusetts Lowell, Lowell, MA, USA; Massachusetts General Hospital, Boston, MA, USA; Brigham and Women’s Hospital, Boston, MA, USA

## Abstract

Sleep disturbances are known to be aggravated with normal aging. Additionally, sleep disruptions have a potentially bidirectional causal relationship with dementia due to neurodegenerative diseases like Alzheimer’s disease. Predictive techniques that can automatically detect cognitive impairment from an individual’s sleep data have broad clinical and biological significance. Here, we present a deep learning approach based on a transformer architecture to predict cognitive status from sleep electroencephalography (EEG) data. This work uses data from *N* = 1, 502 subjects from the Multi-Ethnic Study of Atherosclerosis (MESA) cohort. Our transformer model achieves 70.22% accuracy at the binary classification task for distinguishing cognitively normal and impaired subjects based on their sleep EEG. Our method outperforms traditional feature handcrafting, which has an overall accuracy of 57.61% for the same task. We use a sparse regression model to understand and interpret the information captured by each learned feature from our transformer model. To our knowledge, this is the first effort to use deep learning to predict cognitive impairment from sleep metrics.

## Introduction

Changes in sleep macro- and micro-architecture are a hallmark of healthy aging [12]. Normal aging is known to be associated with many sleep macro-architectural changes, including reductions in total sleep duration, deep sleep time, rapid eye movement (REM) sleep time, and sleep efficiency, and increased compensatory light sleep time, sleep latency, and sleep fragmentation [24]. Sleep micro-architectural changes, including reduction in oscillatory activity such as slow waves and spindles, are also known to occur as people get older. Sleep disruptions are strongly associated not only with normal, age-related cognitive decline but also with dementia due to neurodegenerative diseases such as Alzheimer’s disease (AD) [13, 10]. Epidemiological studies have reported strong associations between reduced self-reported sleep duration and cognitive impairment in the elderly [8]. Research suggests that sleep behavior is intricately linked to amyloid and tau accumulation, the two main neuropathologies implicated in AD [15, 22, 23, 9, 11]. While many interesting open questions remain about potential bidirectional causal connections between sleep and AD, nonetheless sleep metrics have emerged as promising candidates for noninvasive biomarkers for AD [20]. In fact, sleep-based predictors could be used to identify at-risk individuals for cognitive evaluations for recruitment to secondary prevention trials for AD[15]. At the same time, sleep disturbances are considered a modifiable risk factor for AD[6]. There is a need for predictive tools that can automatically detect cognitive impairment from sleep.

While much of the existing research on the links between sleep and cognition relies on simple self-reported [14] or actigraphic [2, 5] sleep duration measures, recent analyses of objective metrics from electroencephalography (EEG) capturing sleep macro- and micro-architecture have provided new insights on how quantitative sleep traits relate to different types of cognitive functions [4]. A key contribution of this study was the identification of 23 (out of over 150) objective sleep metrics that were associated with cognitive performance and processing speed in two large human cohorts: Multi-Ethnic Study of Atherosclerosis (MESA) and Osteoporotic Fractures in Men (MrOS). This analysis, however, was based on a predefined set of handcrafted EEG features.

The data science revolution of the last decade has been fueled by representation learning techniques that implicitly learn suitable features from the data and tend to outperform traditional machine learning techniques that rely on feature handcrafting. Here, we present a deep learning approach to predict cognitive status from sleep EEG data using features that are learned directly from the raw data. We show that a representation learning model based on a transformer architecture [19] which predicts features from the raw EEG time series data is more accurately able to detect cognition than a more traditional model where a large array of precomputed sleep features are input into a fully-connected neural network. Additionally, we use a sparse regression model to understand the information captured by each learned feature from our transformer model. To our knowledge, this is the first effort to use deep learning to predict cognitive impairment from sleep measures. In the subsequent sections, we describe our methodology (including the network architecture, cohort details, and training/validation strategies) and results (model accuracy assessment and interpretation).

## Methods

Sleep architecture is analyzed by examining EEG data, which is typically collected as part of an overnight multimodal sleep study known as polysomnography (PSG). Here, we use sleep EEG and cognitive testing data from an elderly cohort to train and validate a transformer model for our prediction task. We rely on a publicly available dataset for developing and validating the transformer model. Our methodology is described below.

### Data Description

The Multi-Ethnic Study of Atherosclerosis dataset (MESA) is a multi-center longitudinal study that aims to investigate the shift from subclinical to clinical cardiovascular disease. The MESA study included 6,814 asymptomatic men and women of black, white, Hispanic, and Chinese-American ancestry, 2,237 of whom were also participants in the MESA Sleep Study, an ancillary study under MESA. The sleep research participants underwent one full night of unattended PSG. Data were collected across six sites across the United States: Wake Forest University, Columbia University, Johns Hopkins University, University of Minnesota, Northwestern University, and University of California Los Angeles. MESA protocols were approved by the Institutional Review Board at each field center, and all participants gave written informed consent at their respective sites [25, 3].

### PSG

PSG data were collected using the Compumedics®Somte PSG system (Compumedics Ltd., Abbotsford, Australia). The recording montage included cortical EEG, bilateral electrooculography (EOG), chin electromyography (EMG), bipolar electrocardiography (ECG), thoracic and abdominal respiratory inductance plethysmography (RIP) with auto-calibrating inductance bands, airflow measured by nasal-oral thermocouple and nasal pressure cannula, leg movements, and finger pulse oximetry. The data were scored by trained sleep technicians following American Academy of Sleep Medicine (AASM) guidelines to generate epoch-by-epoch sleep labels assigning each 30-s epoch into the five categories wake, REM, N1, N2, and N3, where the last three are non-NREM (NREM) subcategories. The entire cohort’s sleep data were scored at a central Sleep Reading Center. In this study, we utilize raw EEG data and sleep stage labels as inputs to the machine learning model. MESA EEG includes central, frontal, and occipital EEG data sampled at 256 Hz and captured by three channels: Fz-Cz, Cz-Oz, and C4-M1. We utilize the C4-M1 EEG channel as our primary input.

### Handcrafted Sleep Features

For rigorous benchmarking of our feature learning capability, we use a handcrafted feature set as a reference. For handcrafted feature computation, we referred to Djonlagic et al. [4], which generated an exhaustive list of 173 sleep metrics from the MESA dataset. In our analysis, we relied only on objective sleep metrics that are computable from EEG, which reduced the set to a final total of 132 handcrafted features. For a head-to-head comparison between learned and handcrafted features, we separately train a 4-layer fully-connected neural network that receives the 132 handcrafted features as inputs. For model interpretation, we assign these features into 10 broad data-driven domains: 1) total sleep time (TST), 2) sleep efficiency, 3) sleep macro architecture, 4) absolute (slow) power, 5) slow, delta relative power, 6) alpha, sigma, beta power (NREM), 7) alpha, sigma, beta relative power (REM), 8) slow spindles, 9) fast spindles, and 10) spindle frequency.

### Neuropsychological Testing

The Cognitive Abilities Screening Instrument (CASI) is widely used to assess global cognitive function [17]. It is offered in multiple languages and was explicitly developed for cross-cultural studies of dementia. This test includes 25 items from 9 cognitive domains summed to provide an overall cognitive function score on a 0–100 point scale. A score of less than 90 is considered to be indicative of mild cognitive impairment (MCI) [21].

### Data Assembly

Only high-quality PSG data were used in this study. MESA evaluates PSG quality using a 1-7 point scale based on the duration of artifact-free data across the channels (highest quality score: 7, lowest quality score: 1). We used data from all subjects with a PSG quality rating > 6 and a sleep duration of 7.5 hours to ensure data integrity. Additionally, our analysis included only those subjects from whom all 132 handcrafted EEG features were reliably computed as determined by outlier analysis. This led to a total cohort size of *N* = 1, 502. Cohort demographics are provided in Table 1.

**Table 1.**
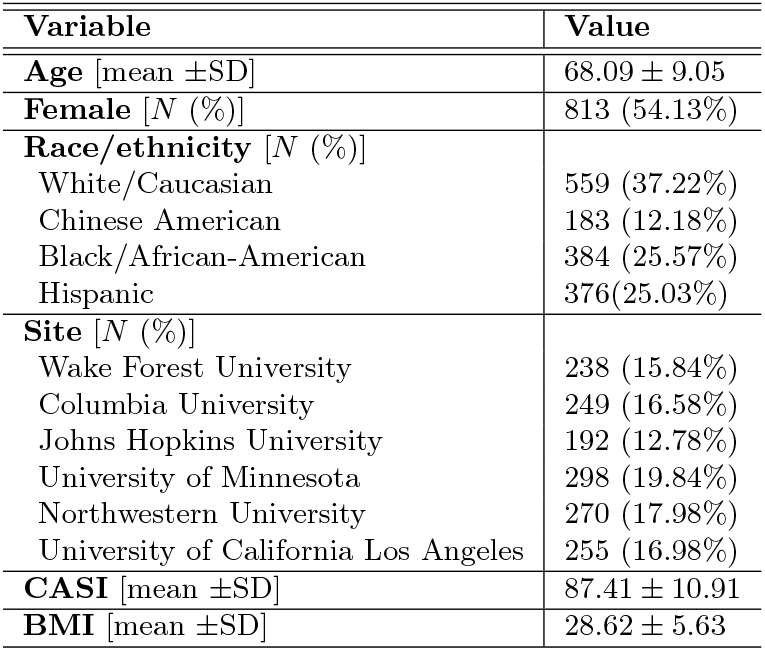
Demographic characteristics of MESA participants with sleep and cognition data used in this study.

### Network Architecture

The deep learning model used in this study is based on the transformer network design with a sequence-to-sequence (Seq2Seq) architecture [16]. A Seq2Seq model typically consists of an encoder and a decoder subnetwork. Because the purpose of our study is to predict a low-dimensional variable (cognition score), we only use the transformer encoder subnetwork. The encoder in our model employs a multi-head attention layer, which is a module for calculating attention for each input feature so that the network can focus on only the most important features during training.

A schematic of the transformer encoder network is depicted in Figure 1. The network receives time series from the whole night as input, with the duration set to span 1200 30-s epochs (10 hours). The transformer encoder employs a constant latent vector of fixed length (denoted *d*) across all of its layers. To keep this length fixed, we utilize a trainable projection module consisting of 3 fully-connected layers to map a 30-s EEG signal window with 7, 680 samples (256 Hz × 30 s) to a d - 1 dimension and then concatenate the output with a PSG sleep stage label value derived from a subject’s hypnogram. After the projection and the concatenation, the input size to the transformer encoder that consists of 2 transformer encoder layers is 1201 × d with a start token, whose state at the output of the transformer encoder combines all the extracted information across all the epochs. Therefore, the output of the second transformer encoder layer at the start position, with size 1 × d, which captures all relevant information, is alone fed into the next level, which is a fully-connected neural network with 4 layers. Age, sex, race, and site – four baseline covariates are included as additional inputs to this fully-connected network which generates the predicted CASI score as its output. In addition, we retrieve attention from the output of the multi-head module in the second transformer encoder layer to determine which transformer features are significant.

**Figure 1.**
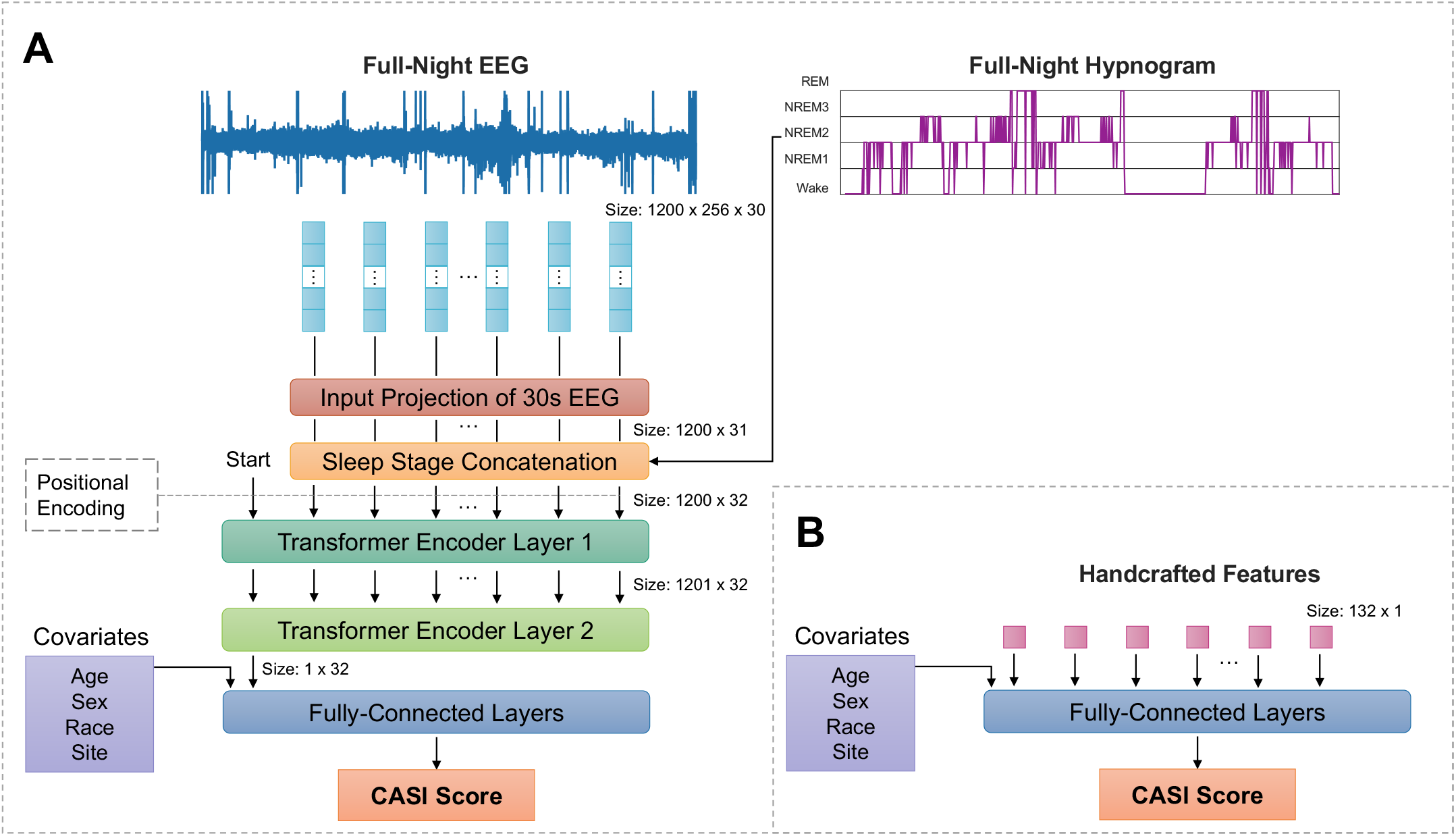
Network overview. (A) The transformer encoder network architecture with raw EEG time-series inputs concatenated with sleep stage labels (wake, REM, N1, N2, N3), which uses two encoder layers for feature extraction. Attention values from 32 features from the transformer encoder layers and four covariates (age, sex, race, and site) are passed into a fully-connected neural network with 4 layers, which generates the predicted CASI cognition score as the final output. (B) A reference neural network architecture with 4 fully-connected layers that receives as inputs a set of 132 handcrafted sleep features (precomputed from the EEG and the sleep labels) and four covariates (age, sex, race, and site) and generates the predicted CASI cognition score as the final output.

### Network Training

The network was implemented and trained on PyTorch using an NVIDIA RTX 3090 graphics card. The hyper-parameters of learning rate and batch size were set to be 0.0001 and 20 respectively. The constant latent vector length *d* was set to 32. The network was trained using an *L*^2^ loss function, and the loss was minimized using the Adam optimization algorithm for 500 iterative epochs [7]. The full dataset comprising *N* = 1, 502 MESA participants was split into a training subset of size 1,042 and an independent validation subset of 460.

### Evaluation Metrics

To evaluate the transformer model, we utilize a set of model performance metrics widely relied upon in the machine learning field to assess classifier performance. The quintessential tool capturing the details of a classification model’s performance is the confusion matrix. Correct and incorrect classification proportions are represented by the diagonal and off-diagonal components, respectively. The accuracy for each class is thus represented by the diagonal members. In addition to providing the confusion matrix, we also compute the sensitivity, specificity, precision, and recall. We also report the *F*_1_ score, which is the harmonic mean of precision and recall. Because it overlooks actual negatives, we note that, despite its popularity, the *F*_1_ score’s effectiveness is limited. We, therefore, also calculate the Matthews Correlation Coefficient (MCC), which is the only metric that captures all of the elements in the confusion matrix in a single scalar value. The MCC computes the correlation between observed and predicted binary classes and hence is symmetric to both positive and negative class definitions.

### Model Interpretation

A key challenge with representation learning models is their “black-box” nature which makes them less intuitive and difficult to interpret. To understand the features generated by the transformer model, we perform feature analysis by comparing the transformer features with the handcrafted feature set via sparse regression. We use Least Absolute Shrinkage and Selection Operator (LASSO), which is a regression approach that combines variable selection and regularization [18]. LASSO aims to preserve a small but most important set of regression coefficients, while setting the rest to zero. In this manner, the LASSO extracts only the most meaningful features in a linear model regression model.

## Results

### Accuracy Comparison

As shown in Figure 2, our transformer encoder network trained using whole-night raw sleep EEG and sleep stage labels has an overall accuracy of 70.22%. The neural network trained using the 132 handcrafted sleep features, on the other hand, has an overall accuracy of 57.61%. The feature learning approach, therefore, outperforms the feature handcrafting approach by a margin of 12.61% in terms of overall accuracy. As shown in Table 2, other evaluation metrics too exhibited sizable margins of improvement for feature learning over feature handcrafting: 14% improvement in sensitivity, 12% in specificity, 12% in precision, 13% in *F*_1_ score, and 25% in MCC. The results are based on 32 transformer features vs. 132 handcrafted features. This suggests that the 32 features are more information-rich and potentially less redundant despite their lower dimensionality than the predefined feature set.

**Figure 2.**
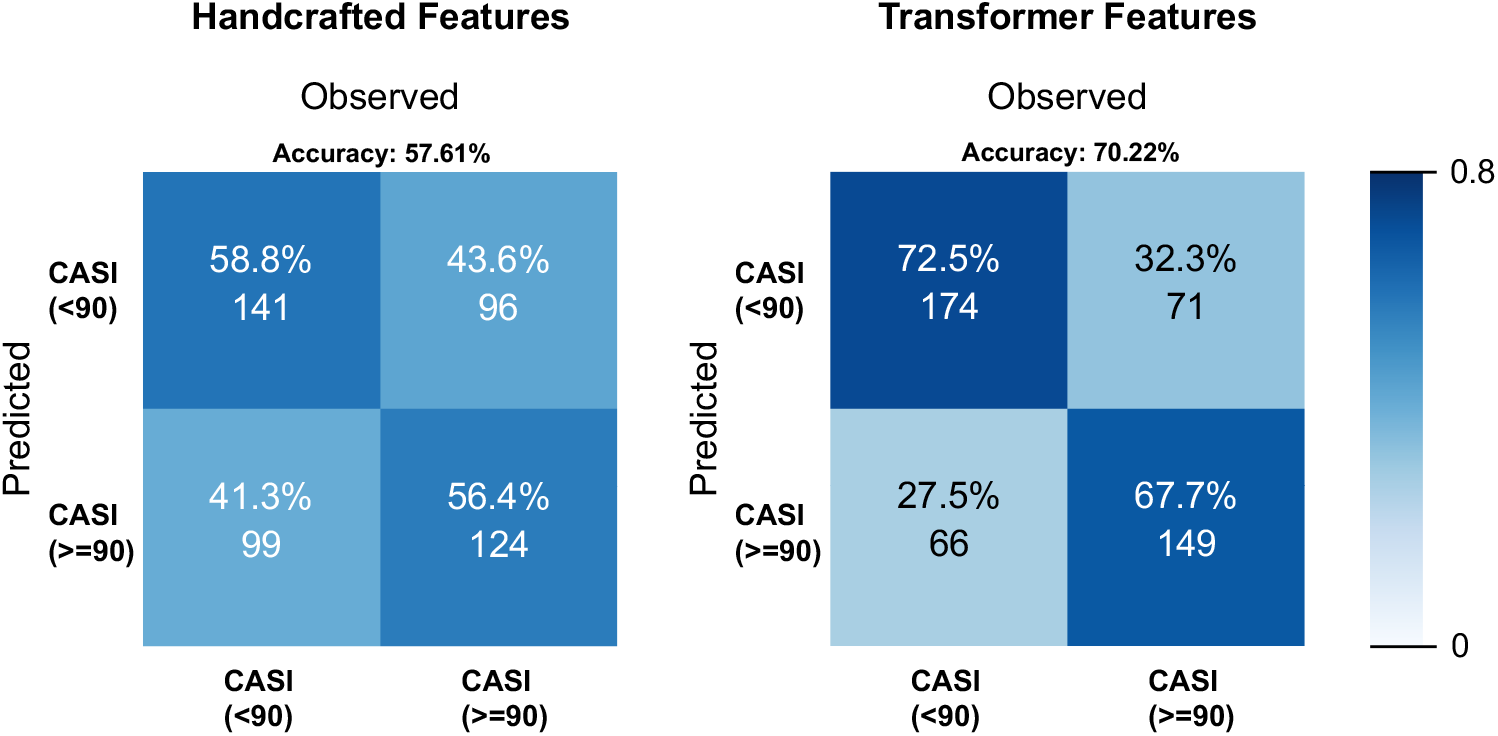
MESA CASI binary classification. Confusion matrices comparing the prediction accuracies of feature handcrafting (overall: 57.61%) and transformer-based feature learning (overall: 70.22%).

**Table 2.** Comparison of classifier performance metrics for feature handcrafting and transformer-based feature learning.

| Metric | Handcrafted Features | Transformer Features |
| --- | --- | --- |
| Sensitivity/Recall | 0.59 | 0.73 |
| Specificity | 0.56 | 0.68 |
| Precision | 0.59 | 0.71 |
| $F_1$ score | 0.59 | 0.72 |
| MCC | 0.15 | 0.40 |

### Transformer Feature Characterization

Figure 3 displays transformer feature ranking based on attention, which is derived by averaging each element from the output of the multi-head layer in the second transformer encoder layer at the start epoch across two subject groups, one with high CASI (above 90) and another with low CASI (below 40). By doing so, we determine which transformer characteristics are statistically relevant to the classifier outcome. We observe that significant features in the high-CASI group receive negative or no attention in the low-CASI group and vice versa. More specifically, transformer encoder features 5, 9, and 22 in the red boxes in Figure 3 receive higher attention in the high-CASI group but receive negative attention in the low-CASI group. Transformer encoder features 3, 12, and 13 in the green boxes, on the other hand, receive higher attention in the low-CASI group but negative attention in the high-CASI group.

**Figure 3.**
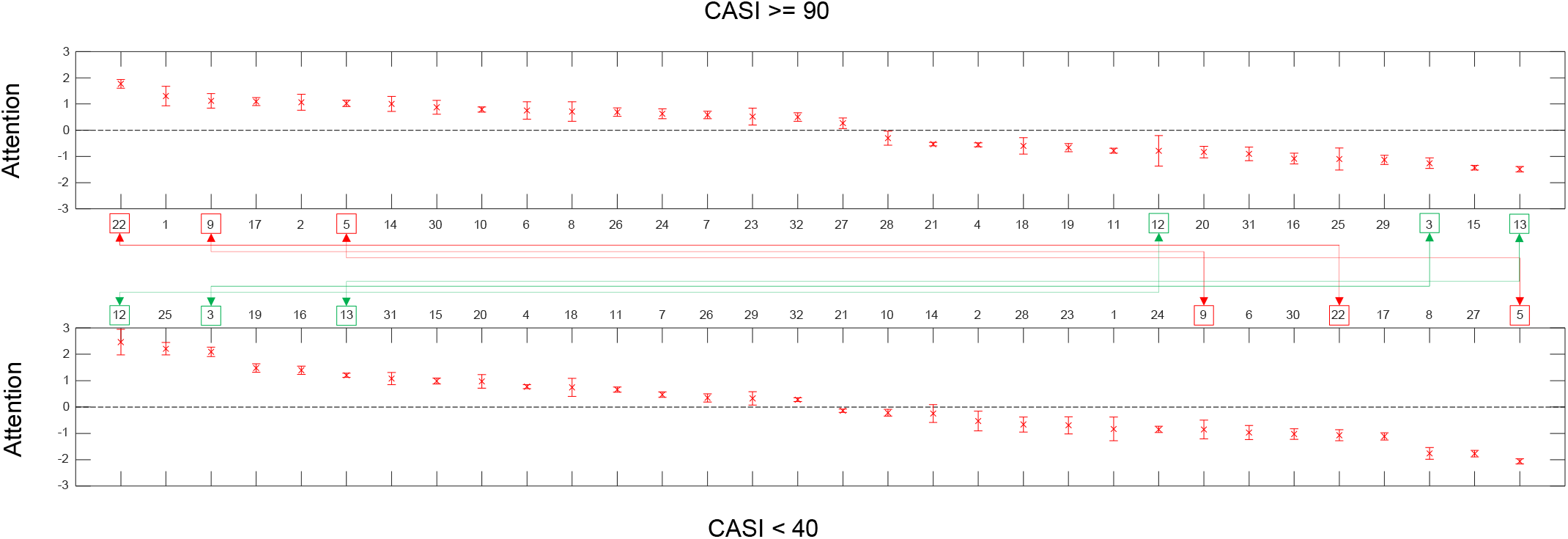
Feature ranking. Transformer feature ranks based on attention for high-CASI (above 90) and low-CASI (below 40) groups. The attention was derived from the second transformer encoder layer’s multi-head attention module at the start point. Features 22, 9, and 5 (located in red boxes) receive more attention from the transformer model in the high-CASI group but less attention in the low-CASI group. Features 12, 3, and 13 (located in green boxes) receive more attention in the low-CASI group but less attention in the high-CASI group.

### Transformer Feature Interpretation

To understand and interpret the information content captured by each transformer feature, we computed sparse regression coefficients mapping the transformer features to the handcrafted feature set. Figure 4 displays a summary of the information content of all 32 features. For this visualization, we accumulated the LASSO coefficients into 10 domains capturing different facets. These domains are based on a data-driven grouping of all handcrafted features as described in Djonlagic et al. [4].

**Figure 4.**
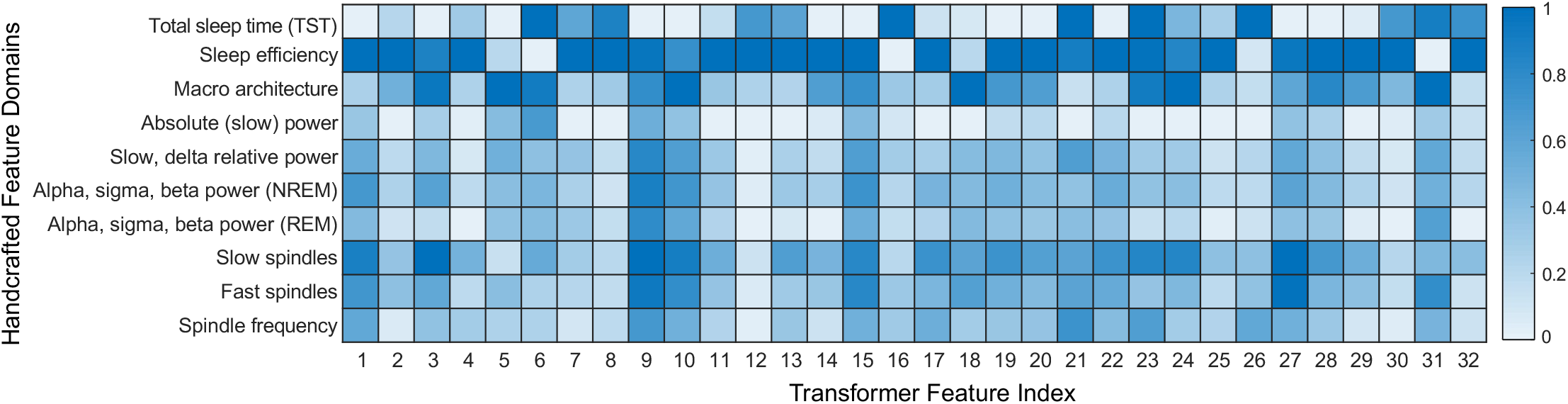
Transformer feature interpretation. LASSO coefficients for the 32 transformer features grouped into 10 broad domains.

For the low-CASI group, 4 out of the top 7 transformer features (12, 13, 16, 31), which contain more total sleep time (TST) information, receive more attention from the model. 5 out of the top 7 transformer features (3, 12, 13, 19, 25) contain sleep efficiency information. 3 out of the top 7 transformer features (3, 19, 31) contain macro-architecture information. 3 out of the top 7 transformer features (3, 13, 19) capture slow spindles. 3 out of the top 7 transformer features (3, 19, 31) contain fast spindles information. Lastly, only 1 out of the top 7 transformer features (31) contain slow, delta relative power information.

For the high-CASI group, the top 7 transformer features (1, 2, 5, 9, 14, 17, 22), which contain less TST information, receive more attention from the model. 6 out of the top 7 transformer features (1, 2, 9, 14, 17, 22) contain sleep efficiency information. 5 out of the top 7 transformer features (1, 9, 14, 17, 22) contain slow spindles information. 4 out of the top 7 transformer features (2, 5, 9, 14) contain macro-architecture information. 4 out of the top 7 transformer features (1, 5, 9, 22) contain slow, delta relative power information. 3 out of the top 7 transformer features (1, 9, 22) contain fast spindles information.

## Discussion

We have presented a machine learning framework for predicting a person’s cognitive status from their full-night sleep EEG data. While the predictive value of sleep metrics has been studied before, to our knowledge, this is the first attempt to build a machine learning model for mapping an individual’s sleep (as quantified by PSG) to their cognition (measured using the CASI score). We show that our feature learning approach, which relies on a transformer model, outperforms traditional feature handcrafting by a sizable margin in terms of an array of performance evaluation metrics.

Although, the MESA dataset used in this study has good representation for mildly impaired individuals (CASI<90), very low CASI scores (CASI<60) are under-represented. This leads to some data imbalance that may affect the underlying model which is set up for a regression task. Owing to this limitation, we report only binary classification results for this model. The threshold of 90 is consistent with a clinical MCI diagnosis based on CASI. Since there is good overall representation of each binary class for this threshold, the model’s performance for this classification task is robust.

In our current setup, we use attention from the second transformer encoder layer at the start point which combines information from all the epochs from the first encoder layer. As future work, we plan to utilize the first encoder layer to see which epochs are important and receive more attention. This will allow us to go further and examine which features each of those epochs carry and thus conduct an epoch-by-epoch interpretation.

One limitation of this study is that it is based on only one night of data from each subject. Since humans tend to have a high degree of night-to-night variability in their sleep patterns, a single night’s data may not be able to capture a person’s sleep habits. Given the cost and complexity of PSG studies, these are rarely conducted for more than one or two nights for a single subject. Improved sleep monitoring accuracy for easy-to-use wearable EEG devices [1] could enable multi-night data acquisition, which, in turn, could produce a longer-term and more complete picture of a person’s sleep habits. Such longer-term assessments may be more meaningful for sleep-cognition mapping efforts such as ours and represent a significant future research direction related to this topic.

## Acknowledgments

This work was supported in part by the NIH grant R21 AG068890 (to J.D., R.S., and S.M.P.). The Multi-Ethnic Study of Atherosclerosis (MESA) Sleep Ancillary study was funded by NIH-NHLBI Association of Sleep Disorders with Cardiovascular Health Across Ethnic Groups (R01 HL098433). MESA is supported by NHLBI funded contracts HHSN268201500003I, N01-HC-95159, N01-HC-95160, N01-HC-95161, N01-HC-95162, N01-HC-95163, N01-HC-95164, N01-HC-95165, N01-HC-95166, N01-HC-95167, N01-HC-95168, and N01-HC-95169 from the National Heart, Lung, and Blood Institute, and by cooperative agreements UL1-TR-000040, UL1-TR-001079, and UL1-TR-001420 funded by NCATS.

